# Pro-inflammatory alveolar macrophages associated with allograft dysfunction after lung transplantation

**DOI:** 10.1101/2021.03.03.433654

**Authors:** Sajad Moshkelgosha, Gavin Wilson, Allen Duong, Tallulah Andrews, Gregory Berra, Benjamin Renaud-Picard, Shaf Keshavjee, Tereza Martinu, Sonya MacParland, Jonathan Yeung, Stephen Juvet

## Abstract

**Purpose:** Lung transplant (LT) recipients experience episodes of immune-mediated acute lung allograft dysfunction (ALAD). We have applied single-cell RNA sequencing (scRNAseq) to bronchoalveolar lavage (BAL) cells of stable and ALAD patients to determine key cellular elements in dysfunctional lung allografts. Our particular focus here is on studying alveolar macrophages (AMs) as scRNAseq enables us to elucidate their heterogeneity and possible association with ALAD where our knowledge from cytometry-based assays is very limited.

**Methods:** Fresh bronchoalveolar lavage (BAL) cells from 6 LT patients, 3 with stable lung function (3044 ± 1519 cells) and 3 undergoing an episode of ALAD (2593 ± 904 cells) were used for scRNAseq. R Bioconductor and Seurat were used to perform QC, dimensionality reduction, annotation, pathway analysis, and trajectory. Donor and recipient deconvolution was performed using single nucleotide variations.

**Results:** Our data revealed that AMs are highly heterogeneous (12 transcriptionally distinct subsets in stable). We identified two AM subsets uniquely represented in ALAD. Based on pathway analysis and the top differentially expressed genes in BAL we annotated them as pro-inflammatory interferon-stimulated genes (ISG) and metallothioneins-mediated inflammatory (MT). Pseudotime analysis suggested that ISG AMs represent an earlier stage of differentiation which may suggest them as monocyte drive macrophages. Our functional analysis on an independent set of BAL samples shows that ALAD samples have significantly higher expression of CXCL10, a marker of ISG AM, as we as higher secretion of pro-inflammatory cytokines. Single nucleotide variation calling algorithm has allowed us to identify macrophages of donor origin and demonstrated that donor AMs are lost with time post-transplant.

**Conclusion:** Using scRNAseq, we observed AMs heterogeneity and identified specific subsets that may be associated with allograft dysfunction. Further exploration with scRNAseq will shed light on LT immunobiology and the role of AMs in allograft injury and dysfunction.

## Introduction

Lung transplantation (LT) is a life-saving treatment option for advanced lung diseases (*1*). However, long-term survival after LT remains significantly lower than after transplantation of other solid organs, despite a substantial improvement in recent years (*2*). The main factor limiting long-term survival is chronic lung allograft dysfunction (CLAD), a progressive fibrotic process that destroys the lung allograft and is driven by inflammation and alloimmunity. Primary graft dysfunction, acute cellular rejection, antibody-mediated rejection and infection are among the main risk factors for CLAD, but how these insults drive CLAD development remains unknown, because the cellular mechanisms that link these risk factors to organ fibrosis remain incompletely understood.

Bronchoalveolar lavage (BAL), in which saline is instilled into the lung via bronchoscopy and then collected using suction, allows sampling of immune cells in the distal pulmonary compartment. In the past two decades, there have been a number of cellular phenotyping studies focusing on associations between clinically defined entities (e.g. acute cellular rejection and CLAD) and limited sets of predetermined cell subsets in the BAL (reviewed in (*3, 4*)), including neutrophils (*5*) and lymphocytes (*6*). However, classifying BAL cells on the basis of a limited set of predefined cell membrane-associated proteins may not uncover the degree of heterogeneity present in the samples nor reveal the contributions of cell populations not classifiable using the markers employed. In addition to that, flow cytometric characterization of alveolar macrophages (AMs) – the most abundant cell population in the BAL – is significantly hindered by their autofluorescence.

AMs play a critical role in maintaining pulmonary homeostasis through interactions with the alveolar epithelium, and during acute inflammation by orchestrating pro-inflammatory and profibrotic responses through phagocytosis and secretion of inflammatory cytokines and reparative molecules (*7–10*). Historically, AMs and other macrophages have been categorized as either M1 macrophages, which respond to bacterial or viral infections, or M2 macrophages, which respond to parasitic infections (*11*). Since M2 features have been observed during wound healing, they are also often described as anti-inflammatory macrophages (*12*). More recent evidence, however, suggests a much more complex and dynamic programming of macrophages that does not conform to this simple dichotomy (*13*). Further, the ontogeny of AMs differs significantly between conditions of homeostasis and inflammation. Recent murine studies suggest that most tissue resident AMs arise during embryogenesis and self-maintain with a minimum contribution from peripheral monocytes (*14–16*). In contrast, a single-cell RNA sequencing (scRNAseq) study of BAL cells from sex-mismatched LT recipients showed that donor AMs in the human lung are replaced by recipient monocyte-derived macrophages (MoMs) (*17*). The use of scRNAseq to study AMs has the potential to deepen our knowledge of this cell population – which was previously considered to be relatively uniform – by allowing us to uncover important heterogeneity within it.

In recent years, scRNAseq has been applied to the study of lung cells including AM, both in humans and in animal models of homeostasis and disease (*17–27*). Two recent studies of BAL cells in patients with COVID-19 infection have focused on tracking viral RNA in infected cells (*28*) and studying immune responses to the disease (*29*). Both reports revealed substantial differences between AMs from patients with mild disease compared to those with illness. However, to our knowledge, no study has specifically examined the association between disease severity and AM transcriptional states on a single cell level.

Our goal is to identify novel cellular contributors to the loss of lung allograft function and CLAD. Here, we have applied scRNAseq to BAL cells from LT recipients with stable lung function or acute lung allograft dysfunction (ALAD) to test the hypothesis that distinct and pathogenetically important AM transcriptional states might emerge during lung allograft dysfunction. We uncovered a landscape of 14 gene expression programs with evidence for functional specialization of AM in the BAL samples of LT recipients. Moreover, our data suggests that there might be associations between allograft dysfunction and two distinct clusters that are only present in the BAL samples of LT recipient with acutely impaired lung function.

## Materials and Methods

### Ethics statement

The study was approved by University Health Network Research Ethics Board (15-9531). All participants provided written informed consent for sample collection and subsequent analyses.

### Patient selection

Six (one female and five male) LT recipients were selected, three with acute lung allograft dysfunction (ALAD, defined as a decrease in the forced expiratory volume in one second [FEV_1_] by 10% or more from the maximum of the two preceding FEV_1_ measurements) and three with stable lung function. We excluded patients if the cause of ALAD was suspected to be infection, identified by focal opacities on chest imaging and/or mucopurulent secretions at bronchoscopy. The demographic characteristics of the study participants are provided in Table 1.

**Table 1.** Clinical characteristics of stable and ALAD patients.

|  | Stable (n=3) | ALAD (n=3) |
| --- | --- | --- |
| Recipient age at LTx (mean) | 63.7±4.7 | 53.7±22.5 |
| Male n (%) | 3 (100%) | 2 (66.6%) |
| D/R sex mismatch | 1 (33.3%) | 2 (66.6%) |
| Native lung disease |  |  |
| Pulmonary Fibrosis/ Interstitial lung disease | 3 | 0 |
| COPD/Emphysema | 0 | 2 |
| Cystic Fibrosis | 0 | 1 |
| CMV D/R |  |  |
| D+/R- | 1 | 1 |
| D+/R+, D-/R+ | 2 | 1 |
| D-/R- | 0 | 1 |
| Culture microbiology |  |  |
| Commensal flora | 3 | 2 |
| No growth | 0 | 1 |
| Previously treated acute cellular rejection | 1 (33.3%) | 0 |
| Respiratory symptoms at time of bronchoscopy | 0 | 1 (33.3%) |
| FEV1 at time of bronchoscopy, median (IQR) | 1.9 (1.9, 2.4) | 2.1 (1.7, 2.99) |
| Baseline FEV1, median (IQR) | 2.1 (2.09, 2.38) | 2.62 (1.95, 4.31) |
| Time from LTx to bronchoscopy (days) | 545.7±181.3 | 821.7±775.9 |
| Acute rejection grade |  |  |
| A0 | 3 | 2 |
| AX | 0 | 1 |
| B-grade |  |  |
| B0 | 1 | 0 |
| BX | 2 | 3 |

### BAL sample collection

Patients underwent bronchoscopy for surveillance or for diagnosis of ALAD. BAL was obtained according to a standard protocol in accordance with international guidelines (*2*). Two 50 mL-aliquots of saline were instilled with the bronchoscope in a wedged position in either the right middle lobe or lingula. Aspiration of BAL fluid was performed after each aliquot was instilled. Fresh BAL samples were transferred to the research laboratory on ice within 10 minutes of sample acquisition.

### Isolation of BAL cells

BAL samples were centrifuged at 100*g* for 5 minutes at 4°C to pellet the larger and more fragile epithelial cells. The supernatant was then transferred to a new tube for a second centrifugation at 400*g* for 5 minutes at 4°C to pellet the remaining cells. The two cell pellets were combined by resuspending in 500 ul of cold PBS + 0.5% fetal bovine serum. Cells were counted and transported to the UHN genomics center for library preparation within 30 minutes of BAL sample acquisition.

### Capture, library construction and sequencing

A total of 11 μl of the single cell suspension and 40 μl barcoded Gel Beads were loaded onto a Chromium Chip A (10x Genomics) to generate single-cell gel bead-in-emulsion. Single cell capture and downstream library construction were performed using 10x Genomics 3’ expression V2 chemistry. Full-length cDNA along with cell-barcode identifiers were PCR-amplified and sequencing libraries were prepared. The constructed library was frozen for later sequencing on the Illumina^®^ NovaSeq platform.

### scRNAseq data alignment and quality control

Cell Ranger (*30*) was used to perform sample de-multiplexing, barcode processing and single-cell 3’ UMI counting with human GRCh38 as the reference genome. Specifically, splicing-aware aligner STAR was used for FASTQ alignment. Cell barcodes were then determined based on distribution of UMI count automatically. Each individual sample was run through Seurat v3 (*31*) with the following filtration criteria applied to each cell: gene number between 200 and 5000, UMI count between 1000 and 30,000 and mitochondrial gene percentage below 10%. Prior to downstream analysis, we also removed genes for ribosomal proteins (those whose names started with RPL or RPS) as suggested by Mould and associates (*18*). After filtration, the remaining cells (stable, 3044 ± 1519 cells; ALAD, 2593 ± 904 cells) were subjected to further analysis.

### Cell annotation

Cells were normalized using the “log normalization” method in Seurat v3 and run through the SingleR package (*26*) to annotate each individual cell in comparison to the Human Primary Cell Atlas (*32*). Cellular annotation was added to the Seurat object as meta-data. In the Seurat object of each individual sample, cells annotated as “Macrophage” were selected for further analysis.

### scRNAseq integration, dimensionality reduction and clustering

The 3 stable and 3 ALAD samples were downsampled to 1000 Macrophage cells per sample. Then the 3 stable samples were merged and integrated using Seurat SCTransform (*31*). A similar procedure was applied to the 3 ALAD samples. For each of the integrated datasets (3 stable BAL and 3 ALAD BAL) the following analyses have been performed: Principal component analysis (PCA) was performed using the top 3,000 most variable genes. Then t-distributed stochastic neighbour embedding (tSNE) was performed on the top 30 principal components for visualization of the cells. Subsequently, a graph-based clustering approach using K-nearest neighbor analysis was performed on the top 15 principal components with a resolution of 0.8 for the granularity of the downstream clustering.

### DGE analysis

To measure differential gene expression (DGE) for each cluster, MAST (*33*) in Seurat v3 was used to perform differential analysis using normalized data. Similarly DESeq2 (*34*) was performed for comparing raw RNA expression between individual samples as well as between integrated stable and integrated ALAD samples.

### Cell-cell interaction

We employed the CellChat R package (*35*) to infer the intercellular communication based on normalized scRNAseq data. CellChat predicts major signaling inputs and outputs by comparing the cell interaction in the human CellChat Database – which includes cell contact, extracellular matrix-receptor, and secreted signaling data from KEGG and the literature.

### Comparison of stable and ALAD datasets

In order to compare the integrated stable and ALAD datasets, we applied the results from DGE analysis and Seurat to the ClusterMap algorithm (*36*). ClusterMap matched clusters across the two datasets and provided ‘similarity’ as a metric to quantify the quality of the match.

### Pathway analysis

Gene ontology (GO) and Gene Set Enrichment Analysis (GSEA) were performed with ClusterProfiler (*37*) which supports statistical analysis and visualization of functional profiles for genes and gene clusters. In GSEA analysis, 50 hallmark gene sets in MSigDB were used for annotation.

### Genotyping scRNAseq samples using a single nucleotide variation (SNV) calling algorithm

To identify donor and recipient cells from scRNA-seq data we used a previously published algorithm, Vireo (*38*), which was designed to identify cells derived from genetically distinct individuals in synthetically pooled samples. Since transplant samples contain a mixture of cells from genetically distinct individuals, we reasoned that Vireo would enable us to deconvolute donor and recipient cells. Cell Ranger (*30*) alignments were piled up with the requirement that a called SNV be covered by at least 15 unique cell barcodes, at least 10 unique barcodes supporting the alternative allele, an allele fraction of at least 1%, and to have at least one read with a supporting mismatch outside of the first and last 5 base pairs of the read. We inferred the strand for each potential SNV by requiring that at least 90% of the reads covering the SNV be derived from the same strand. We further filtered these SNVs to remove A>G RNA edits as these are non-genic and not useful for genotype assignments. We identified an SNV as an A>G edit if it was a strand-specific A>G change and present in REDIportal (*39*) or overlapped an Alu element according to RepeatMasker annotations downloaded from UCSC (*40*). Genotypes were called by running Vireo v0.3.2 with two genotypes and our SNV dataset with SNVs removed.

After genotyping, we initially observed that there were low-quality genotype calls. Therefore, we developed a second filtering step to remove cells that appeared to have poor genotype assignments. First, we filtered the potential SNVs that were expressed in less than 10 percent of the assigned cells in either genotype. Next, we calculated the summary allele fraction for each genotype/SNV combination summing the (total alternative allele counts) / (total alternative + reference allele counts). Next, for each genotype we selected SNVs that were predictive of the cell’s genotype by requiring SNVs to have a receiver operator curve (ROC) AUC >= 0.5 (AUC calculated as |AUC – 0.5| * 2). Next, we selected for SNVs that are homozygous in one genotype and reference in the other as heterozygous SNVs are noisier due to the sparsity of scRNAseq data and ambient RNA contamination. We selected these by requiring the absolute value of the difference between overall allele fractions between the two genotypes be greater than or equal to 0.85. The liberal cut-off below 1.0 is to account for ambient RNA contamination. Next, for each cell we selected the SNVs that are covered by at least one molecule and calculated the average of the absolute value of the difference of each SNV minus the summary SNV for each genotype. The expectation is that each cell should have a low score compared to the assigned genotype and a high score compared to the other genotype. Doublets and unassigned cells are expected to have scores around 0.5 as they are a mix of both genotypes. Finally, we calculated a threshold for each genotype to remove low quality cells with scores higher than the cut-off. The cut-off was set by averaging the difference scores from the cells assigned to the given genotype + 4 median absolute deviations (limited the cut-off to be between 0.15 and 3). We selected all the predicted singlets for each genotype that fell below the cut-off for further analyses.

### Identifying a minimum set of SNVs to visualize cell genotype assignments

To find a minimum set of SNVs for each cell we first filtered the list of SNVs to have an AUC of 0.75 (see above) and a mean difference of 0.4 (to include heterogeneous SNVs). Next, we applied a greedy algorithm to find a set of SNVs that covers genotype 1 by selecting each SNV that covers the largest number of uncovered cells from both genotypes. We repeated this for the second genotype to find SNVs that are present in both genotypes.

### Functional analysis, flow cytometry, and cytokine array

Fresh BAL samples from another 3 ALAD and 4 stable LT recipients were collected and cells were obtained by centrifugation at 400*g* for 5 minutes at 4°C. BAL cells were cultured at one million cells per millilitre of media (DMEM with 10% FBS) with or without 20ng/ml of LPS. Cultured cells and supernatant were harvested after 16-18 hours and cryopreserved for subsequent experiments. Cells were fixed using 1% formaldehyde before staining with the following surface markers: anti-CD45-BV605 (clone: HI30, BD Biosciences), anti-CD11b-PerCP/Cy5.5 (clone: M1/70), anti-CD163-PE/Dazzle 594 (clone: GHI/61), anti-HLA-DR-Alexa Fluor 647 (clone: L243), and anti-CD32-PE-Cy7 (clone: FUN-2) (all from Biolegend, San Diego, CA), and anti-CD206-APC-Vio 770 (clone: DCN228) (Miltenyi Biotec, Germany). Cells were washed, permeabilized (Cytofix/Cytoperm, BD Biosciences), and intracellular staining was performed with anti-CXCL10-PE (clone: J034D6, Biolegend) and anti-CD68-BV785 (clone: Y1/82A, Biolegned). Cell data was acquired on a BD LSRII flow cytometer and analyzed using Flowjo software. Media supernatants were used to measure released cytokines by BAL cells using a 13-plex bead-based immunoassay and a human macrophage LEGENDplex (Biolegend, San Diego, CA).

## Results

### Structure of the LT BAL cellular compartment at single cell resolution

Here, we present the single cell transcriptional landscape of BAL samples from LT recipients who were either stable (n=3) or experiencing ALAD (n=3) at different time points post-LT (Fig 1A and Table 1). Single cell transcriptomes from each sample were analyzed separately, following standard quality control protocols (Fig S1). We applied a reference-based annotation algorithm (ClusterR) to identify the main cell populations (Fig 1B) as well as a combination of t-distributed stochastic neighbor embedding (tSNE) and shared nearest neighbor clustering to identify robust cell clusters (Fig 1C). Analysis of the top 5 differentially expressed genes in each cluster (Fig 1D) showed that the samples were heterogeneous, including epithelial, endothelial and immune cells; the majority of the cells were AM. We then analyzed cell-cell interactions using the CellChat algorithm (*35*). We assessed the total interaction strength of the aggregated cell-cell communication network (Fig 1E), as well as the signaling pathways through which different clusters are predicted to be interacting (Fig 1F). Our data suggested that most interactions between AMs and T cells are through CCL and CXCL pathways, while the interaction pathways between different AM subpopulations are more diverse. After studying these aspects for stable BAL samples, we performed a similar analysis on ALAD samples. Given the predominance of AM over other cell populations in BAL samples, we next examined these cells in greater depth.

**Fig 1.**
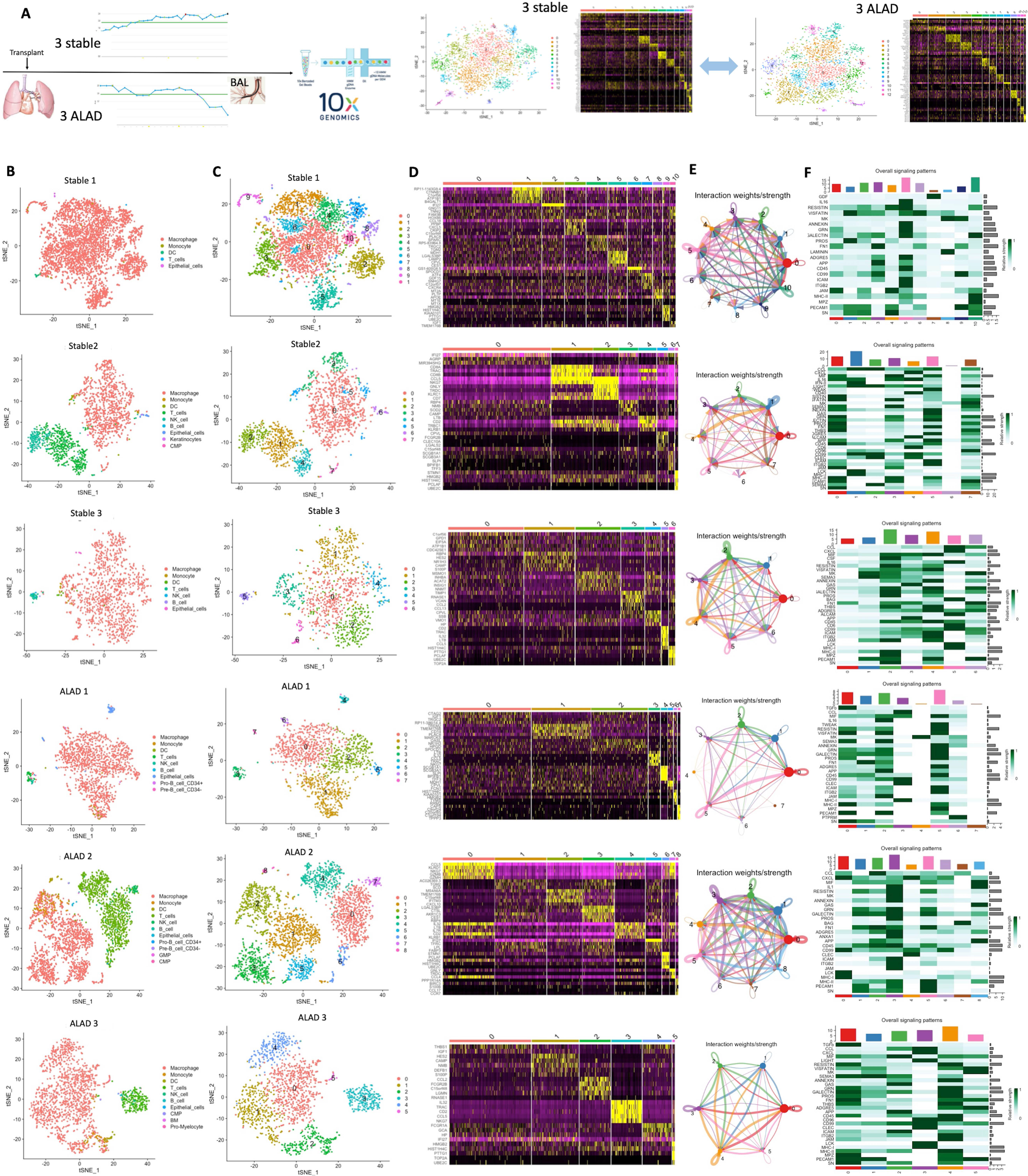
The cellular composition of BAL from LT recipients is revealed by single cell RNA sequencing. **(A)** Study design. Three stable and three ALAD patients with no evidence of pulmonary infection underwent bronchoscopy at varying times post-transplant and BAL was subjected to scRNAseq on the 10x genomics platform. **(B)** Data analysis of individual samples of stable and ALAD LT (one sample at each raw) with reference-based annotation using SingleR algorithm. **(C)** tSNE plot demonstrating clusters of each sample demonstrated between 6 and 10 distinct clusters. **(D)** Distinct gene expression programs depicted in a heatmap of the top differentially expressed genes. **(E)** The total interaction strength (weights) between any two cell groups using circle plots based on the top differentially expressed genes compared to validated molecular interactions using the CellChat algorithm. **(F)** Overall signaling patterns for each cluster are shown in heatmaps where dark green represents a higher relative strength of signal.

### AMs from patients with stable lung function fall into at least 12 clusters

The total number of AMs in the 3 stable BAL samples ranged from 1769 to 4681. We randomly selected a 1000-cell subset (downsample) of AMs from each stable sample to optimize the outcome of integration. Next, we used SCTransform (*31*) to integrate scRNAseq data from all three stable samples (Fig S2A) and data from all three ALAD samples (Fig S2B) to minimize potential batch effects arising from different sequencing runs. Expression of known macrophage transcripts confirmed that the selected cells are AMs (Fig S3). Clustering of the cells identified 12 distinct subsets (Fig 2A) with different patterns of gene expression (Fig 2B). Further analysis revealed that all 13 clusters were represented in each sample (Fig 2C). Our cell-cell communication analysis also demonstrated that each cluster has a distinct signaling network, some with a broader set of pathways (such as clusters 6 and 8) while others appear to use fewer pathways (for example, cluster 9), to interact with other AM clusters (Fig 2D).

**Fig 2.**
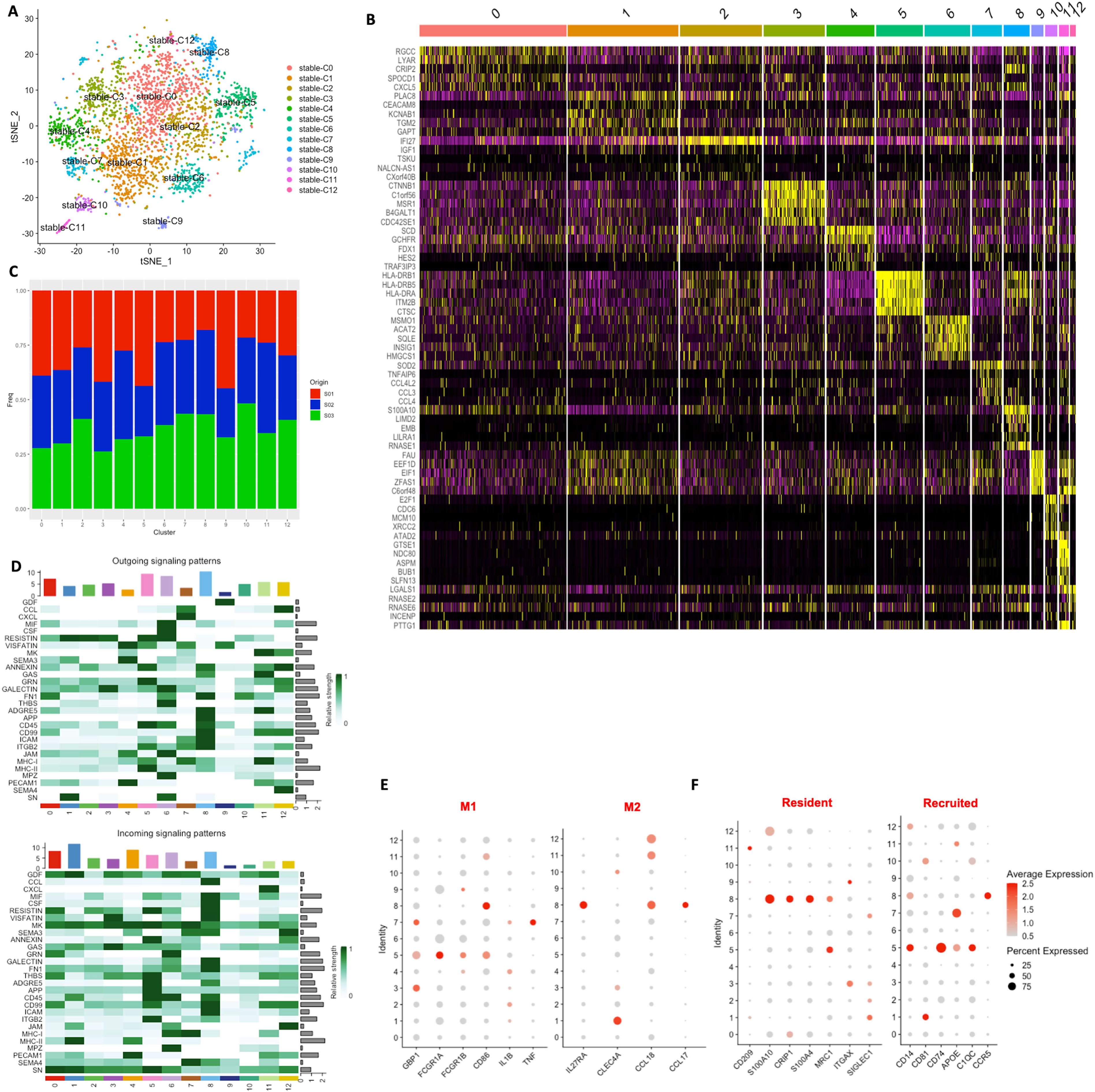
The AM compartment of stable LT recipients exhibits a diversity of transcriptional states. **(A)** tSNE plot of AMs from n=3 stable patients reveals 13 distinct cellular clusters. **(B)** Heatmap depicting the top differentially expressed genes in each of the 13 clusters. **(C)** Frequency histogram showing representation of each cluster in each of the 3 samples. **(D)** Heatmaps demonstrating outgoing (top) and incoming (bottom) signaling patterns for each cluster. **(E)** Expression of canonical M1- (left) and M2- (right) associated genes in AMs from stable patients. **(F)** Expression of genes associated with resident (left) and recruited) AMs. In E and F, circle colour reflects average relative gene expression within the cluster, while the size of each circle reflects the percentage of cells within the cluster expressing the indicated gene.

Next, we examined whether one or more of the identified AM clusters can be classified as M1- or M2-polarized macrophages based on known genes (*41*). In the BAL samples from stable LT recipients, our data reveal that cluster 5 differentially expressed 4 genes associated with M1 polarization while cluster 8 differentially expressed 3 genes associated with M2 polarization (Fig 2E). Overall, the data indicate that expression of canonical M1- and M2-associated genes was distributed over multiple cell clusters, rather than being strictly associated with specific clusters (Fig 2E). Since fetal yolk sac-derived and monocyte-derived macrophages (recruited) can contribute to the AM compartment, we further examined differential expression of genes associated with these populations (*18*). Interestingly, the M1-like cluster 5 differentially expressed *CD14*, *CD74*, *APOE* and *C1QC* – which are associated with monocyte-derived macrophage populations. The M2-like cluster 8, by contrast, most strongly expressed *S100A10, CRIP1, S100A4,* and *MRC1* which are associated with tissue-resident yolk sac-derived AM (Fig 2F). Taken together, these observations suggest that clusters 5 and 8 express genes classically associated with M1-like monocyte-derived AMs and M2-like resident AMs, respectively; however, this summation does not adequately capture the heterogeneity present in AMs after LT.

### Pathway analysis suggests functional specialization for each cluster of AMs

For each of the 12 clusters identified using differentially expressed genes in stable patients, we next performed pathway analysis using GO and GSEA (Fig 3A). Also, we plotted the most differentially expressed genes in each cluster that were also expressed in the majority of the cells of that cluster (Fig 3B). In addition, we studied the incoming and outgoing ligand/receptor signaling network for each cluster (Fig 3C and Fig S4) to gain insight into how the clusters may interact with each other. Further, we examined the cell cycle phase of all AMs in the sample (Fig 3D). We then integrated our findings with data reported in the literature to annotate each cluster based on either a putative function or cycling status (Fig 3E). We identified 3 clusters with specialized anti-microbial functions (PLAC8^+^: extracellular pathogen defense; IFI27^+^: viral defense; CTNNB1^+^: intracellular bacterial defense) (*42–44*), two clusters with specialized immune regulatory functions (SOD2^+^: anti-inflammatory, CD32b^+^: resident Ig-regulated), one cluster with sterol synthesizing and regulating function (MSMO1^+^)(*45*), one cluster with potential scavenger function (SCD^+^) (*46*), one cluster with potential healing and/or profibrotic genes (FN1^+^)(*47*), one cluster with a higher expression of inflammatory markers (CTSC^+^: M1-like, recruited, inflammatory), one cluster with several eukaryotic initiation factors (EIF) (*48*) and two AM populations in the cell cycle (*21, 49*), one with G2/M phase transcripts and another with S phase transcripts. These data suggest that the AM clusters we have identified may have distinct functional properties, some of which have been described previously (Table 2).

**Fig 3.**
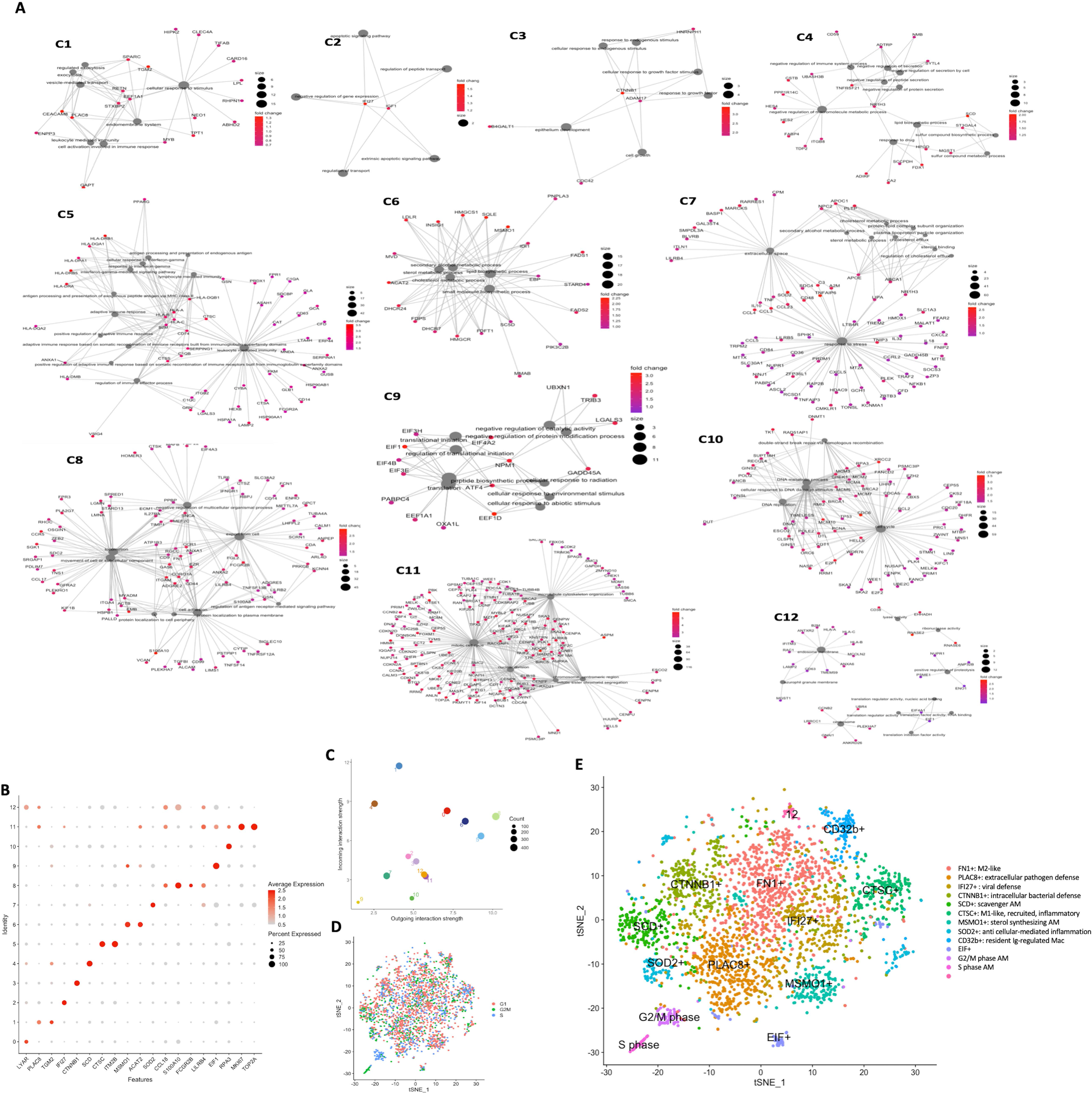
Pathway analysis suggests functional specialization of AMs from stable patients. **(A)** Pathway analysis performed using GO and GSEA revealed distinct gene programs in the 13 clusters. **(B)** Expression of the top differentially expressed genes in each stable AM cluster (x-axis) according to cluster (y-axis). Circle colour reflects average expression within the cluster, while the size of each circle reflects the percentage of cells within the cluster expressing the indicated gene. **(C)** Scatterplot showing total incoming (y-axis) and outgoing (x-axis) ligand/receptor interaction networks for each stable AM cluster. **(D)** tSNE plot of AMs from stable BAL with cell cycle stage indicated. Distinct clusters of G2M-phase (cluster 10) and S-phase (cluster 11) AMs are seen at lower left. **(E)** Functional annotation of AMs in stable BAL samples based on pathway analysis.

**Table 2.** Functional annotation of AMs in stable and ALAD BAL samples

| Stable AM cluster | ALAD AM cluster | Cluster Id | Cluster specificity/function | Previous similar observations | Reference |
| --- | --- | --- | --- | --- | --- |
| stable-C0 | ALAD-C1 | FN1 | M2-like AM | Several top genes (inc. FN1) is found in M2-like cells | Jablonski et al., 2015 (47) |
| stable-C1 | ALAD-C0 | PLAC8 | Extracellular pathogen defense AM | Functional importance shown in mouse models | Ledford et al., 2007 (42) |
| stable-C2 | ALAD-C8 | IFI27 | Viral defense AM | Activated in viral infection via TLR7 activation | Tang et al., 2017 (43) |
| stable-C3 | ALAD-C9 | CTNNB1 | Intracellular bacterial defense AM | β-catenin-mediated bacterial elimination mechanism | Fu et al., 2017 (44) |
| stable-C4 | ALAD-C4 | SCD | Scavenger AM | There is high similarity between this cluster top DGEs and previously reported FABP4 expressing macrophages. | Coulombe et al., 2014 (46) |
| stable-C5 | ALAD-C3 | CTSC | M1-like, recruited, inflammatory AM |  |  |
| stable-C6 | ALAD-C7 | MSMO1 | Sterol synthesizing AM | Lipid metabolism gene signature | Poczobutt et al., 2016 (45) |
| stable-C7 | ALAD-C11 | SOD2 | Anti-cell-mediated inflammation AM |  |  |
| stable-C8 | ALAD-C6 | CD32 | Resident Ig-regulated AM |  |  |
| stable-C9 | ALAD-C2 | EIF | EIF+ AM | mTORC1 signaling pathway | Pugliese et al., 2017 (48) |
| stable-C10 | — | G2/M Phase | G2/M-phase AM | Cycling macrophages | Travaglini et al., 2020 (49);<br>Zilionis et al., 2019 (21) |
| stable-C11 | ALAD-C10 | S phase | S-phase AM | — |  |
| — | ALAD-C5 | ISGs | Inflammatory AM | Similar gene expression observed in AM of severe COVID-19 patients | Liao et al., 2020 (29) |
| — | ALAD-C12 | MTs | Metallothionein-expressing AM | Similar gene expression observed in AM of severe COVID-19 patients | Liao et al., 2020 (29) |

### Similarities and differences between AM subsets in stable and ALAD samples

We next sought to compare the transcriptomes of AMs from LT patients with stable lung function to those from LT patients with ALAD. We analyzed 1000 randomly selected AMs from the ALAD BAL samples using the same approach taken for the stable BAL samples. We observed a similar degree of heterogeneity among ALAD AMs (Fig 4A and Fig S5), with 13 distinct clusters. However, review of the top differentially expressed genes and pathway analysis (Fig S6) demonstrated that these clusters do not in all cases correspond to those observed in samples from stable LT patients. To identify similarities and differences between AM subsets in the two groups, we used Clustermap to perform DGE analysis of each cluster systematically. This approach revealed that there are 11 AM subsets in common between the two groups of patients, and two unique clusters only present in patients with ALAD (Fig 4B and Fig S7A). By comparing the frequency of equivalent pairs of clusters in stable and ALAD samples, we noted that there were five clusters in the ALAD samples which are either absent (two clusters) or present in lower relative frequency (>20% differences between two groups, three clusters) in stable LT recipient BAL (Fig 4C). In addition, despite the fact that some ALAD AMs were in G2/M phase (Fig S7B), our data demonstrated that cells in this phase of the cell cycle only formed a distinct cluster in samples from stable patients. This observation suggests that there may be less transcriptional diversity among AMs in stable BAL samples compared to ALAD samples, leading to greater resolution of S and G2/M phase AMs; alternatively, ALAD AMs might be more rapidly cycling (with a correspondingly shorter S phase), decreasing our ability to resolve differences between these phases of the cell cycle.

**Fig 4.**
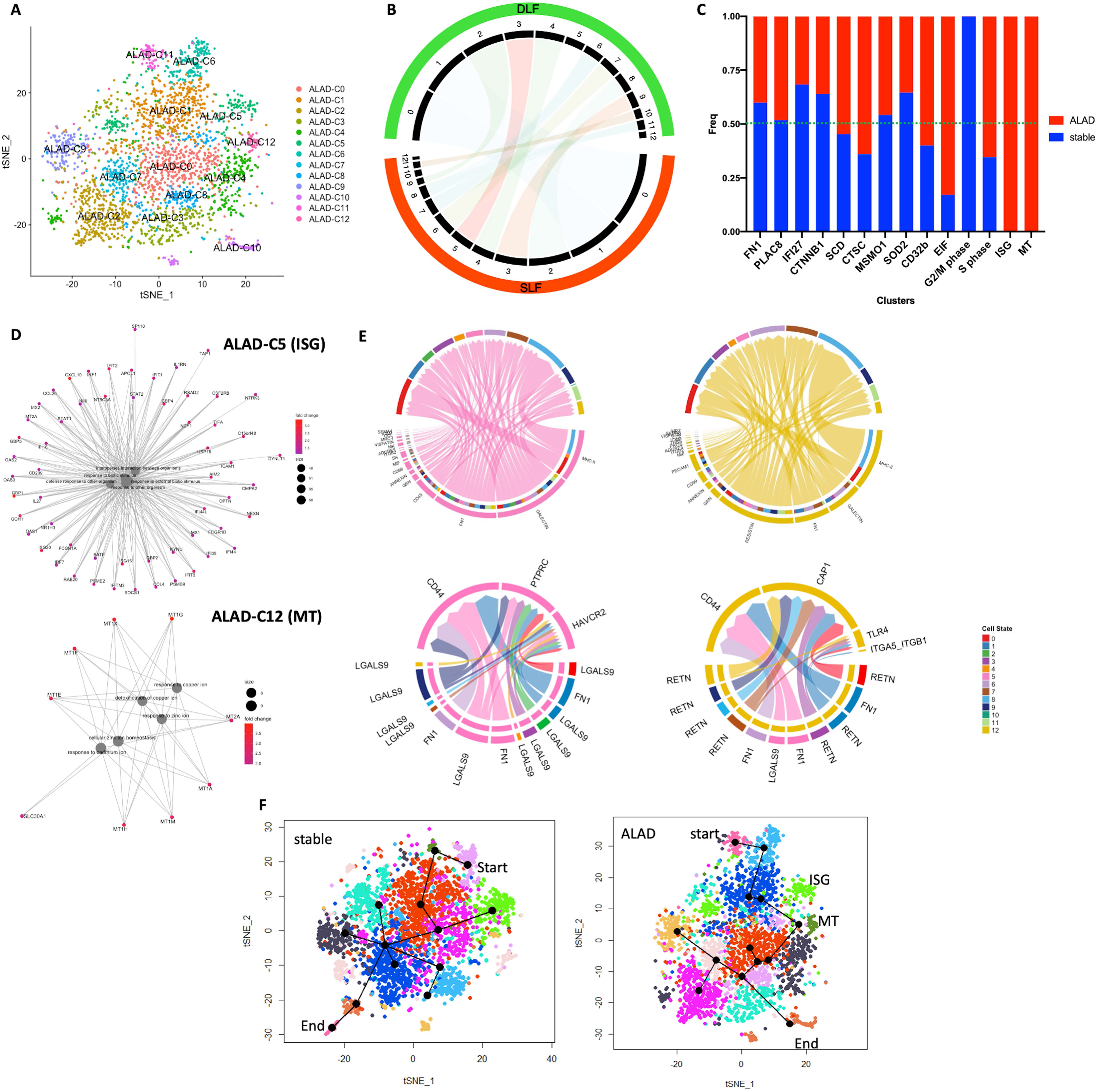
AMs with unique features are present in BAL from patients with acute lung allograft dysfunction. **(A)** tSNE plot of AMs from ALAD patients reveals 13 distinct cell clusters. **(B)** Clustermap analysis comparing ALAD and stable BAL samples reveals 11 shared AM clusters with two clusters uniquely represented in ALAD BAL and one cluster uniquely represented in stable BAL. **(C)** Histogram showing representation of AM clusters in ALAD (red) and stable (blue) samples. Only ALAD samples contain ISG and MT AMs while only stable samples contained AMs forming a distinct G2/M-phase cluster. **(D)** Pathway analysis of ISG and MT AMs from ALAD samples. **(E)** The signaling network of ISG (top left) and MT (top right) AMs and enriched signaling pathways for ISG (bottom left) and MT (bottom right) AMs through which they interact with other ALAD AM clusters. **(F)** Pseudotime trajectory analysis was performed using Slingshot showing similar clusters for start and end point of pseudotime for both stable and ALAD.

We then focused on the two clusters that were unique to ALAD, cell clusters with expression of interferon-stimulated genes (ISGs) and metallothioneins (MTs) to investigate whether they may be involved in allograft dysfunction. Based on the top DGE of ISGs cluster and its pathway analysis, there are several highly expressed transcripts in this cluster that are involved in pro-inflammatory pathways (Fig 4D). Our investigation into the signaling networks through which these two clusters communicate with other AM clusters further support their potential pro-inflammatory role (Fig 4E). MT AMs particularly interact through the RESISTIN-CAP1 pathway, which has been shown to induce a pro-inflammatory response in macrophages (*50*). To further validate our observations about these two sub-populations, we investigated publicly available scRNAseq data from a recent study focusing on the cellular composition of BAL samples from COVID-19 patients (*29*). We identified AM clusters exhibiting a very high degree of similarity to the MT and ISG clusters observed in the ALAD samples (COVID-19 AM clusters 22 and 0, respectively, in Liao et al.). In keeping with the notion that these AM populations may mediate lung inflammation, the majority of AMs in these clusters originated from patients with severe, rather than mild, COVID-19 (Fig S7).

In addition to the clusters unique to ALAD, three additional clusters (EIF, CTSC^+^, and CD32b^+^) were present in higher relative abundance compared to samples from stable LT recipients (Fig 4C). On the other hand, IFI27^+^ and SOD2^+^ cells exhibited a higher relative abundance in stable compared to ALAD samples (>20% differences between two groups, Fig 4C). To further evaluate similarities and differences between AM originating from stable and ALAD patients, we performed pseudotime trajectory analysis using Slingshot (*51*) (Fig 4F). The data revealed that MT and ISG AMs arise at a similar stage in pseudotime, whereas cycling AM were found at the end of the trajectory. In contrast, CD32b^+^ AMs seem to have recently differentiated, perhaps from monocytes. Despite small differences in pseudotime branching, the general trajectory patterns for ALAD and stable samples were similar. In ALAD, MT and ISG AMs were located in the early-intermediate part of the pseudotime trajectory, suggesting that they may have recently differentiated from recruited monocytes in response to specific stimuli.

### Identifying the origin of AMs using SNVs genotype assignments

Work by others (*14, 49, 52, 53*) and data presented here illustrate that the AM compartment contains cycling cells that presumably permit self-renewal. In LT, this raises the question of whether and how long donor AMs persist in the allograft recipient. This question has been addressed by either HLA-A2 or analysis of Y chromosomes in sex-mismatched LT, which have suggested an exceptionally long-term persistence of donor AM (*54, 55*). We chose to examine this issue using an SNV calling algorithm that allowed us to infer the donor and recipient origins of cells – without reference to genomic sequence data – by determining which SNVs were present in the majority of epithelial cells in the sample (Fig 5A). As has been observed previously, our data suggest that there is indeed a replacement of donor-derived by recipient-derived AM in the BAL over time post-transplant (Fig 5B), but in contrast to the prior reports, most of the replacement occurs within the first-year post-transplant. At 3 months post-transplant, most AM were of donor origin; at 12 months post-transplant, only 10% of AM were donor-derived and by 24 months, AM were exclusively recipient-derived (Fig 5C). Furthermore, our data suggest that replacement of AMs are not dependent on particular subsets of AMs and all clusters are eventually replaced by recipient macrophages (Fig 5D).

**Fig 5.**
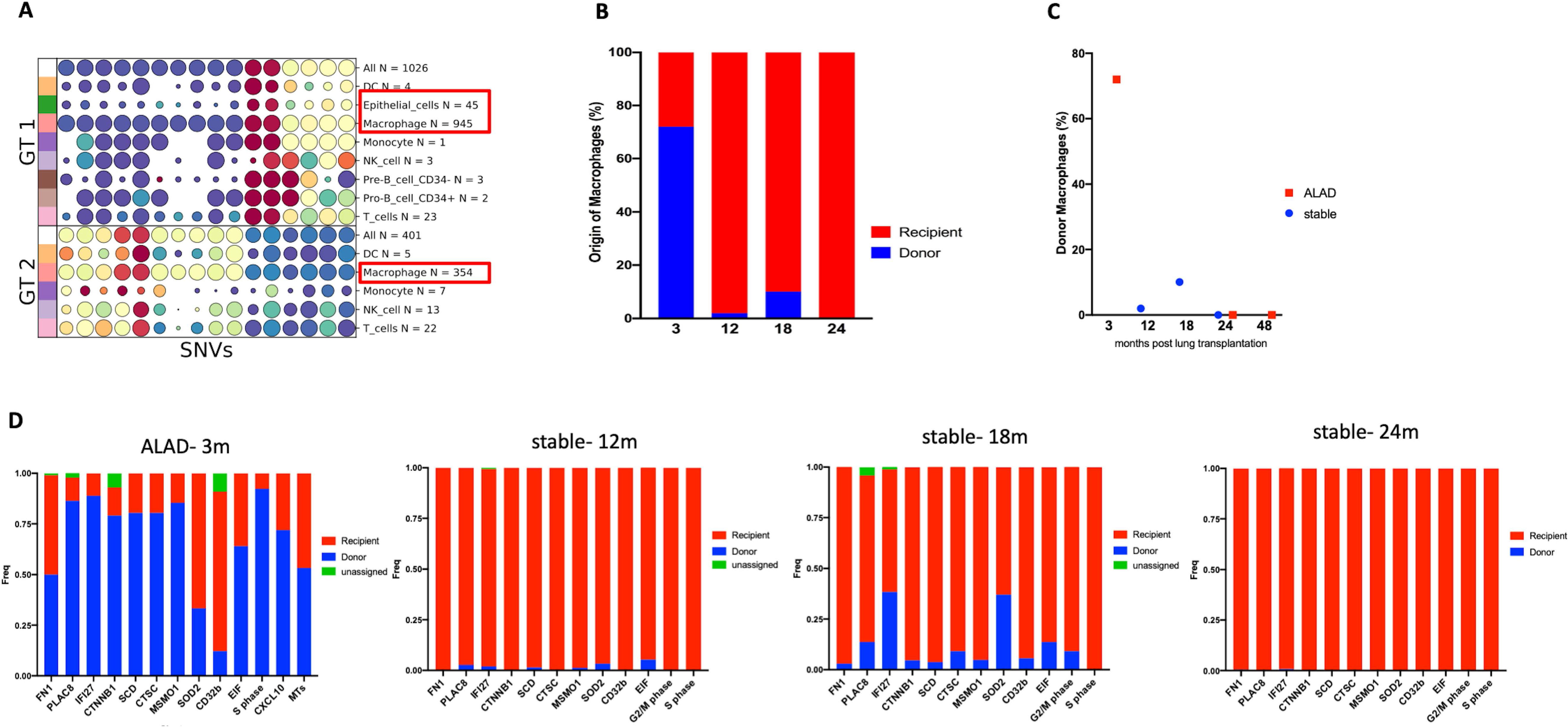
Identification of donor and recipient AMs using an SNV calling algorithm. **(A)** Identification of different SNVs within specific cell populations allowed identification of genotypes 1 and 2 (GT1 and GT2). In this example from an ALAD patient at 3 months post-LT, epithelial cells came from GT1 and are therefore ascribed to the donor. AMs of GT1 outnumbered those of GT2 (red boxes). Circle sizes indicate the proportion of cells in which each SNV is expressed, whereas colours indicate the average minor allele fraction across the cells in each group (red=high, blue=low). **(B)** Histogram showing the proportions of donor (blue) and recipient (red) AMs in BAL samples at various times post-transplant. **(C)** In both ALAD (red) and stable (blue) BAL samples, donor AMs are lost with time post-LT. **(D)** Histogram of AM frequency of origin (donor vs. recipient), of each cluster in 4 stable and ALAD BAL samples showing AMs in all clusters are replaced by recipient cells over time.

### Functional properties of BAL macrophages

To determine whether AMs associated with ALAD could be identified based on cell-associated and secreted proteins, we obtained an independent set of 7 freshly isolated BAL samples (n=4 stable and n=3 ALAD) and stimulated the cells overnight in the presence or absence of LPS. The cells were subjected to flow cytometry to distinguish AM subsets that we identified in scRNAseq data (full gating strategy is presented in Fig S8). To identify ISG AMs, we used CXCL10 in conjunction with CD163. The percentage of CXCL10-expressing AMs was significantly lower in BAL samples from stable patients but, with LPS stimulation, the proportion of CXCL10+ AMs in these samples was increased to the level seen in ALAD samples (Fig 6A). However, there was no further increase in the percentage of CXCL10+ AM in ALAD samples in response to LPS, suggesting there might be a limited number of AM that can acquire this phenotype. Furthermore, we analyzed cytokine and chemokine levels in the AM culture supernatants using a 13-parameter multiplexed cytokine assay. In keeping with the greater presence of CXCL10+ AMs in ALAD samples, our data revealed that IL-6, TNFα, IFNγ and CXCL10 are released in greater quantities by AM from ALAD compared to stable samples (Fig 6B). These findings validate the observation that ISG AMs are associated with ALAD in LT patients.

**Fig 6.**
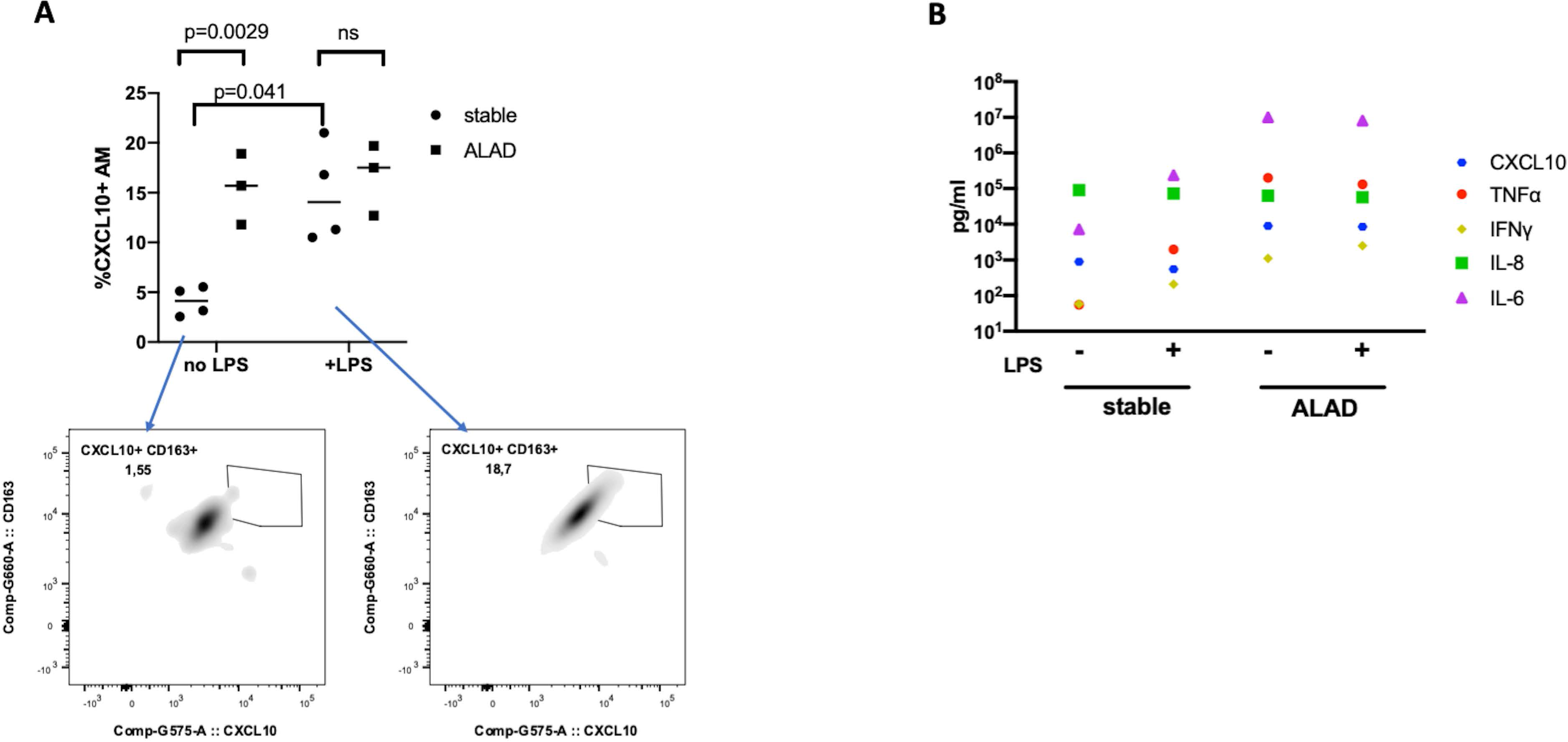
Identification of CXCL10-expressing AMs in ALAD patients. In an independent set of BAL samples (n=4 stable and n=3 ALAD), AMs were cultured overnight in the presence or absence of LPS. AMs were identified as CD68+HLA-DR+ cells. **(A)** By intracellular staining, CXCL10 expression was elevated in AMs from ALAD compared to stable BAL samples; CXCL10 expression was augmented in stable AMs but was not further increased in ALAD AMs. Panels below show representative examples of CXCL10+CD163+ stable AMs without (left) and with (right) LPS stimulation. Repeated measures mixed effects model with Sidak’s post-hoc tests. **(B).** Measurement of cytokine released in culture media by freshly isolated BAL cells from 4 stable and 3 ALAD with and without LPS stimulation. From 12 measured cytokines, IL-6, TNFa, IFNg and CXCL10 are released in greater quantities by AM from ALAD compared to stable samples. Data show the median of cytokine measurements for each group (n=4 stable and n=3 ALAD). Each sample was run in duplicate.

## Discussion

The application of scRNAseq to heterogeneous samples has provided a wealth of information on the composition and transcriptional states of a variety of tissues in health and disease. In many respects, BAL cells are ideal for scRNAseq because they are readily accessible through a minimally-invasive procedure and reflect the inflammatory state of the distal lung compartment. The poor outcome of LT compared with other solid organ transplants makes the application of this technology to BAL and lung tissue samples an appealing approach to identify previously unrecognized contributors to episodes of ALAD, which cumulatively predispose to CLAD (*56–58*).

In this study, we used scRNAseq to compare AM transcriptional states in BAL samples from LT recipients with either stable lung function or ALAD. We chose to focus on AMs because 1) they represent the majority of cells in BAL samples and 2) the specific characteristics of AMs in relation to lung allograft dysfunction have not been investigated in detail. Our data reveal a landscape of 14 gene expression programs in the macrophages of LT recipient BAL samples, and that – with the exception of MTs and ISGs – their proportions were remarkably consistent across all 6 samples and at differing times post-transplant. The data therefore reveal a previously unappreciated diversity in LT AMs, with evidence for functional specialization among the populations. In agreement with previous reports (*21, 59, 60*), our data show that the M1 vs. M2 paradigm inadequately captures AM diversity. Rather, transcripts that are traditionally associated with M1 and M2 macrophages appear to be distributed across the populations, with some expressing both M1- and M2-associated transcripts.

We applied a enhanced SNV calling algorithm to the scRNAseq data, which confirmed previous reports (*54*) describing replacement of donor by recipient AMs following LT. In contrast to those studies, which suggested that donor AM replacement may take several years, our findings indicate that donor AMs are mostly replaced within 12 months post-LT. The previous work distinguished donor and recipient AMs on the basis of allogeneic HLA molecules detected using flow cytometry, whereas our study used SNVs. We believe that this approach, with its coverage of the sequenced transcriptome, more accurately attributes each cell to donor and recipient than does HLA typing, since intact allogeneic HLA molecules can move from the surface of one cell to another via trogocytosis or exosome-mediated transfer (*61*). Our findings will require further validation, but the data suggest that relying on the surface expression of allogeneic HLA molecules might overestimate the duration of persistence of donor AMs. This is an important area for future investigation, since the rate of clearance of donor leukocytes over time may have prognostic implications in LT (*62*).

There is a general consensus among scRNAseq studies on murine lung macrophages that three to five distinct subpopulations exist, with some variation based on the nature of the model (*18, 26, 27*). In contrast, human lung scRNAseq datasets have demonstrated substantial disagreement with the murine studies, with two (*49*) to more than 10 (*21, 29*) subsets identified. We therefore compared our data with macrophage profiling studies (using either bulk or single cell RNA sequencing) in lung samples from human and mice. Of the 14 clusters that we identified, some AM subsets, with similar gene profile signatures and function, have already been reported. For example, proliferating macrophages have been demonstrated in several studies (*21, 49*). However, our identification of AM populations in different phases of the cell cycle in stable patients is an observation that has not been reported previously, to our knowledge. There are a number of possible explanations for this finding. First, either decreased diversity of non-cell cycle-related transcripts – or cell cycle synchronization – among AMs in stable compared to ALAD samples might have resulted in distinct S and G2/M populations. Alternatively, AMs in the ALAD BAL samples may have been cycling more rapidly, which might have caused AMs in one cell cycle phase to retain transcripts from the previous phase, leading to an inability to resolve them into distinct clusters. A third possibility is that both monocyte-derived and yolk sac-derived cells are proliferating in the ALAD samples, and transcriptional differences between these cell types might have obscured the transcriptional distinctions between G2/M and S phases. Finally, it is conceivable that a distinct self-renewing AM population exists in stable samples that is lost in ALAD. A further exploration of these possibilities awaits further study.

We observed remarkable and extensive cell-cell communication networks amongst AMs in both stable and ALAD patients, suggesting the existence of complex feedback and feed-forward loops between the different AM transcriptional states in the bronchoalveolar compartment of the transplanted lung; in contrast, communication between AMs and T cells appeared to be more limited, with predicted pathways primarily restricted to chemokine signalling. These findings will require further study, but they suggest that control of the inflammatory tone of the alveolar space in LT recipients is subject primarily to complex inter-relationships between AMs rather than AM-T cell interactions.

Our data demonstrated substantial differences – despite our small sample size and the high genetic diversity associated with human subjects – between stable and ALAD patients. Most notably, ISG and MT AMs were uniquely represented in ALAD samples. An examination of differentially expressed genes in ISG AMs suggested that different pro-inflammatory cascades have been induced, probably in response to IFNγ, including guanylate-binding proteins (GBPs), cytokines and chemokines (CXCL10, CCL4, and CCL2), in addition to several members of the ISG family. Although the IFNγ response in macrophages is usually associated with infections, it is likely that ISG AMs in our ALAD samples were responding to alloimmune-mediated inflammation since we excluded patients with infection from the study. However, it remains unclear whether this cluster of cells represents a developmentally distinct macrophage population, or whether the activation of an ISG program simply reflects the effects of IFNγ on a susceptible subset of AMs. Since these cells were absent from the stable BAL samples that underwent scRNAseq and only represented in small proportions of the stable samples subjected to flow cytometry, we favour the latter explanation. Importantly, AMs isolated from stable patients produced less CXCL10 – one of the most differentially expressed genes in the ISG cluster – than AMs from ALAD patients. This finding validates the scRNAseq observations at a phenotypic level. Nevertheless, it was possible to elicit CXCL10 expression in stable AMs that was comparable to that of ALAD AMs using LPS, indicating that the AM phenotypes we describe here are dynamic.

Interestingly, the MT AMs, which were also uniquely associated with ALAD in our dataset, have not previously been shown to participate in alloimmunity. MTs are a family of metal-binding proteins that maintain homeostasis of zinc and copper, mitigate heavy metal toxication, and alleviate superoxide stress. They also play a role in inflammatory responses and antimicrobial defenses (*63*). Particularly in macrophages, it has been suggested that MTs are required for a pro-inflammatory response (*64*). Although several publications suggest that MTs participate in inflammation, we are unaware of previous reports of an association between MT-expressing macrophages and allograft dysfunction.

We also observed differences in the relative frequency of other AM clusters between stable and ALAD samples that might have biological relevance. Amongst them are EIF, CTSC^+^, and CD32b^+^ cell clusters. There is evidence that different members of the EIF family are associated with attenuating the anti-inflammatory response in macrophages (*65*) which could conceivably play a role in the development of ALAD. Differences in the relative frequency of CTSC^+^ and CD32b^+^ cell clusters were not as pronounced as the EIF population. Based on our observation that the frequency of CD32+ AMs increased in response to LPS in stable patients but not in ALAD patients, we speculate that this cell population might have immunoregulatory properties. Nevertheless, pathway analysis of both CTSC and CD32b+ clusters suggested that they may alternatively contribute to an inflammatory response mediated by AMs. Further work will be required to understand the functional properties of these AMs.

In summary, we have shown that the AM compartment undergoes specific transcriptional alterations in LT recipients experiencing ALAD, with primarily recipient-origin monocyte-derived ISG and MT AMs emerging. To date, most research on ALAD has focused on lymphocyte-mediated adaptive immune responses. While these are undoubtedly central to lung allograft loss, the appearance of specific AM gene expression programs during ALAD suggests that these cells may contribute actively to the loss of lung function and may therefore represent important mechanistic targets.

## Notes

### Competing Interest Statement

The authors have declared no competing interest.

