## Supplementary figures and images for "Pro-inflammatory alveolar macrophages associated with allograft dysfunction after lung transplantation"

### Supplemental Figures

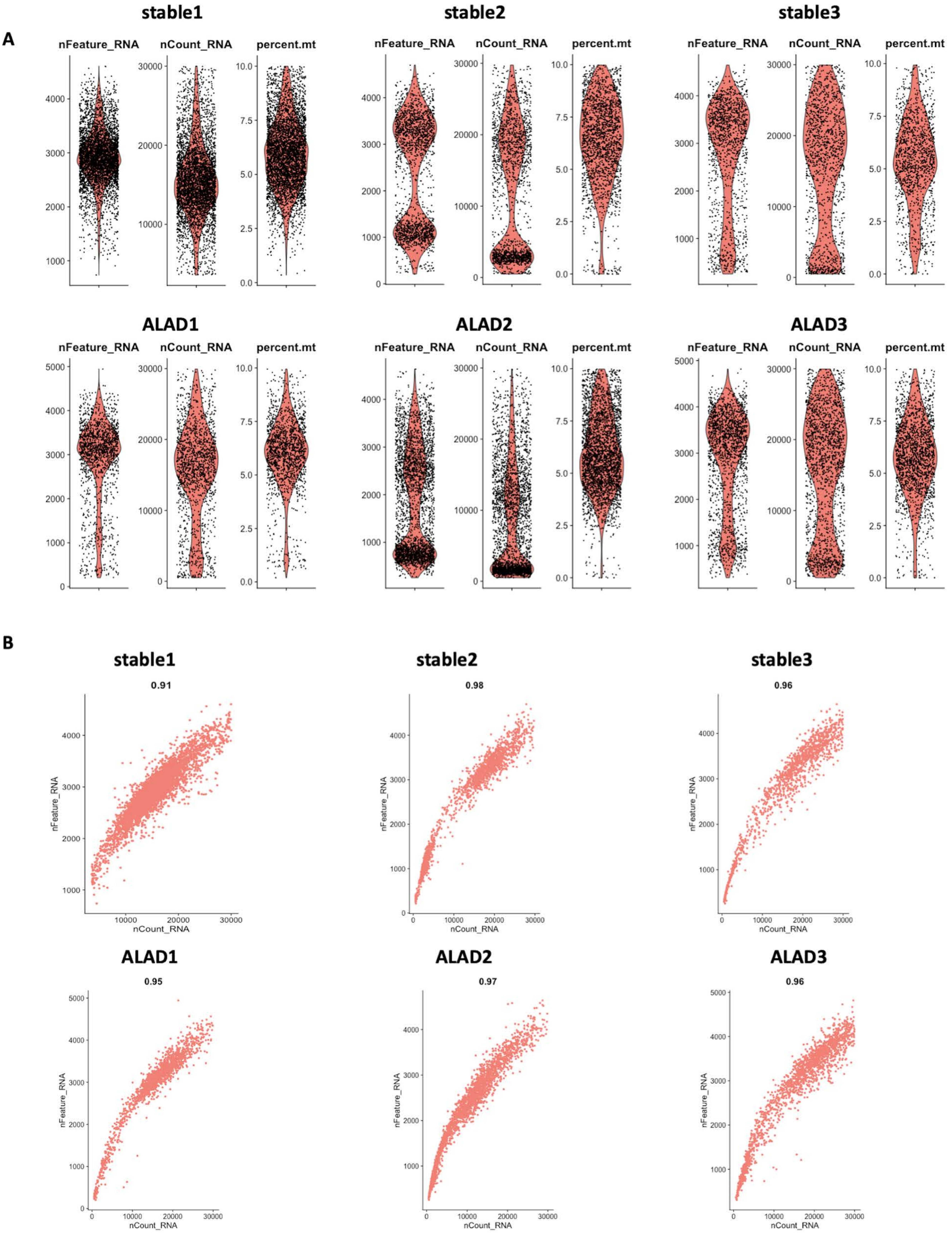

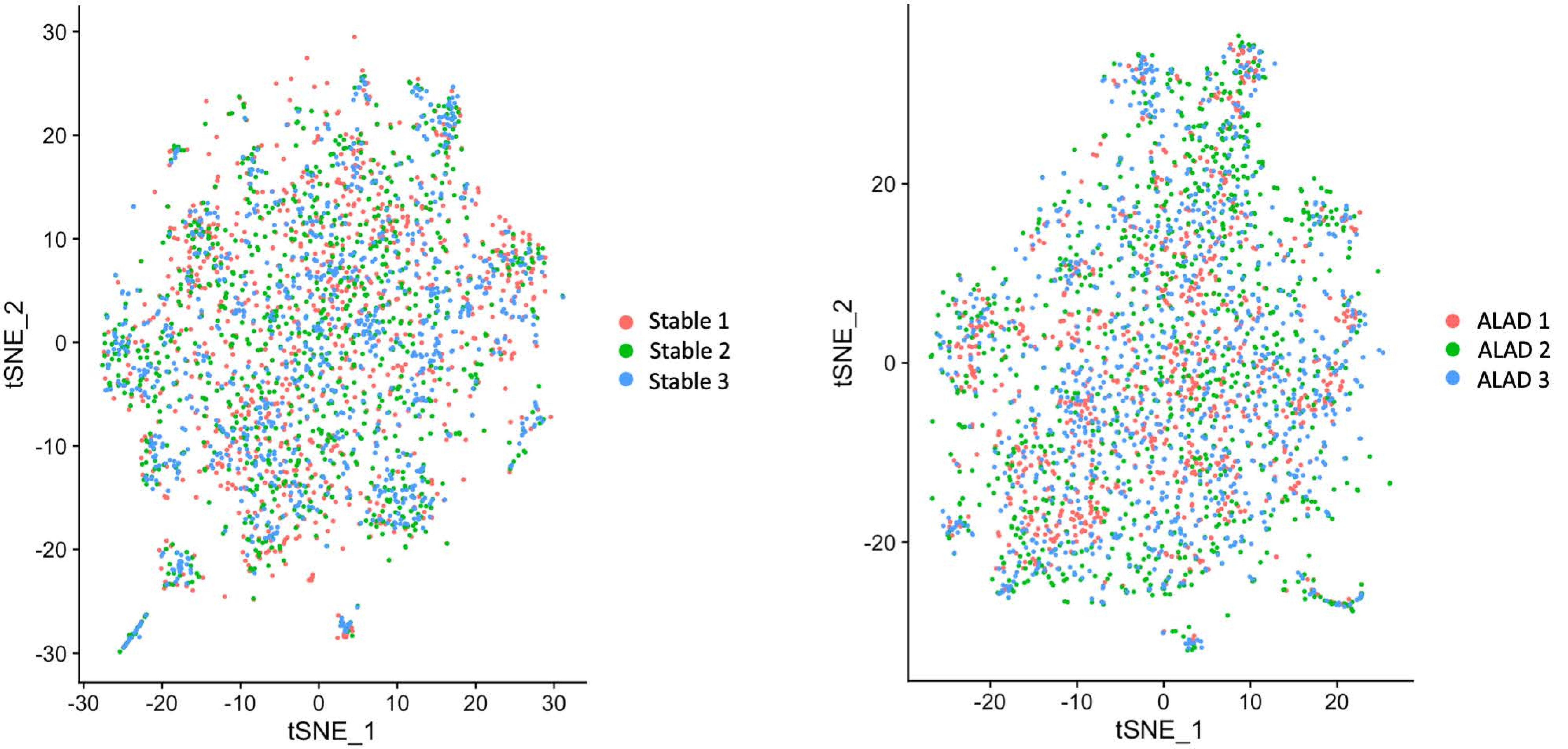

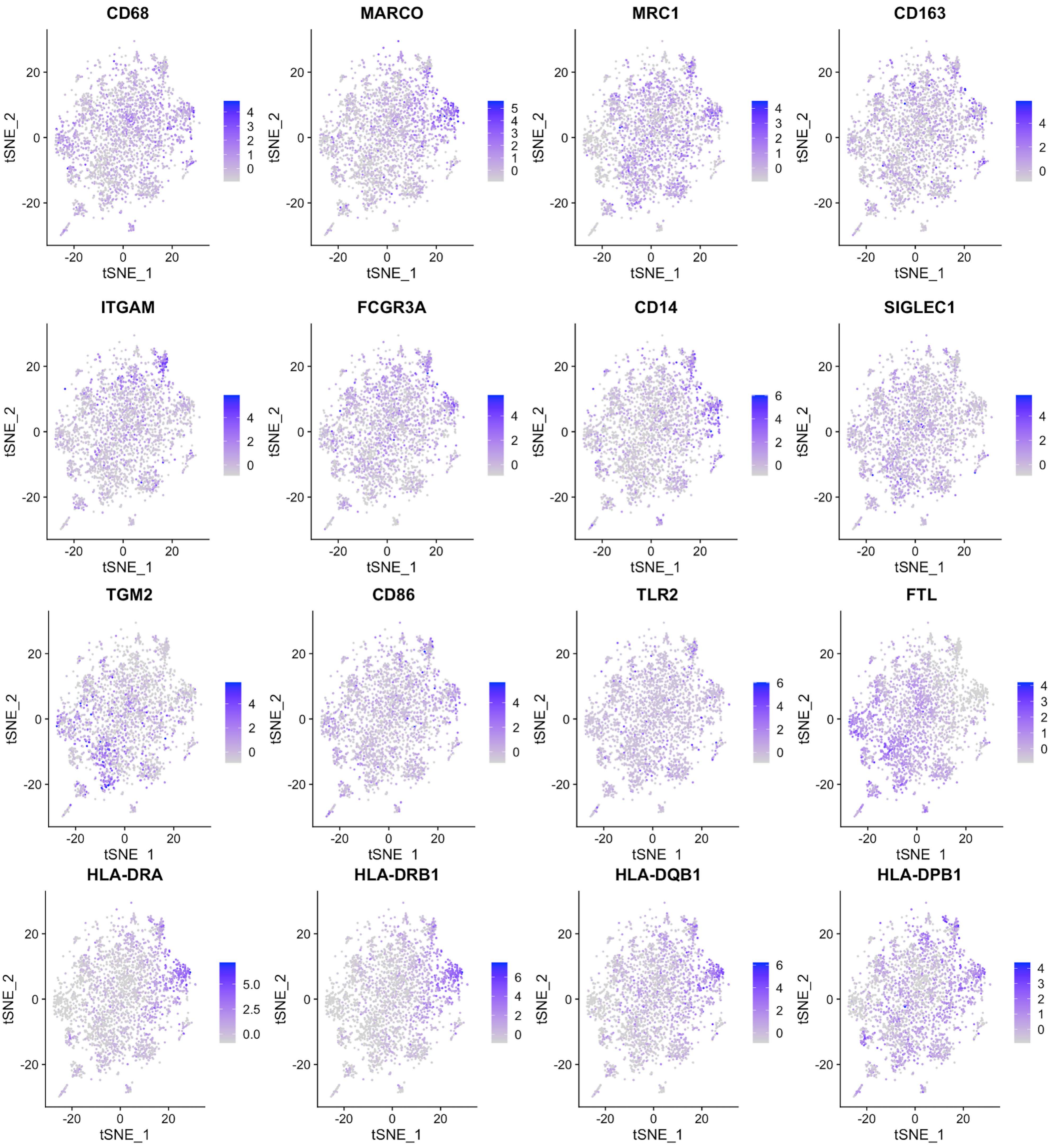

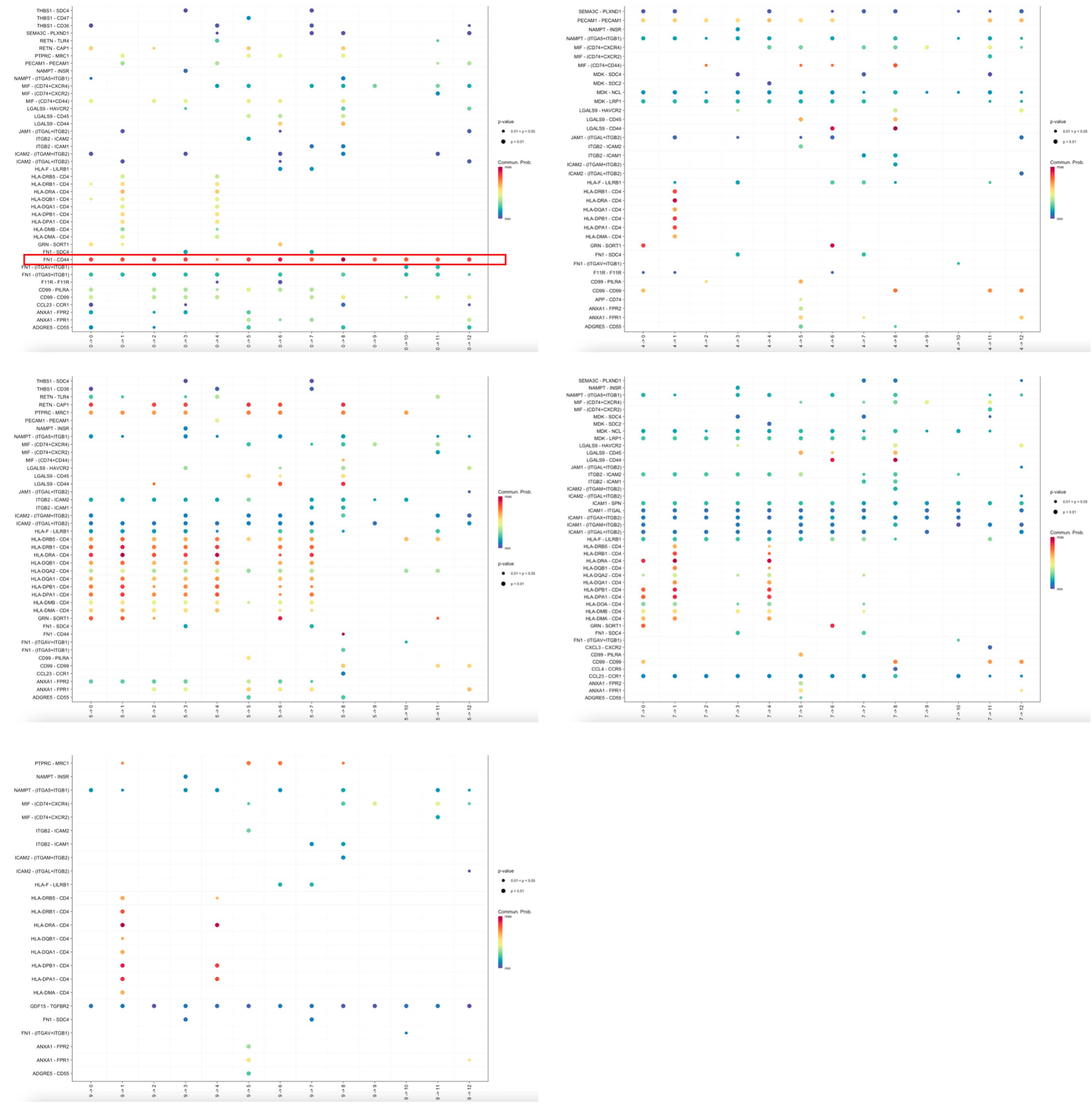

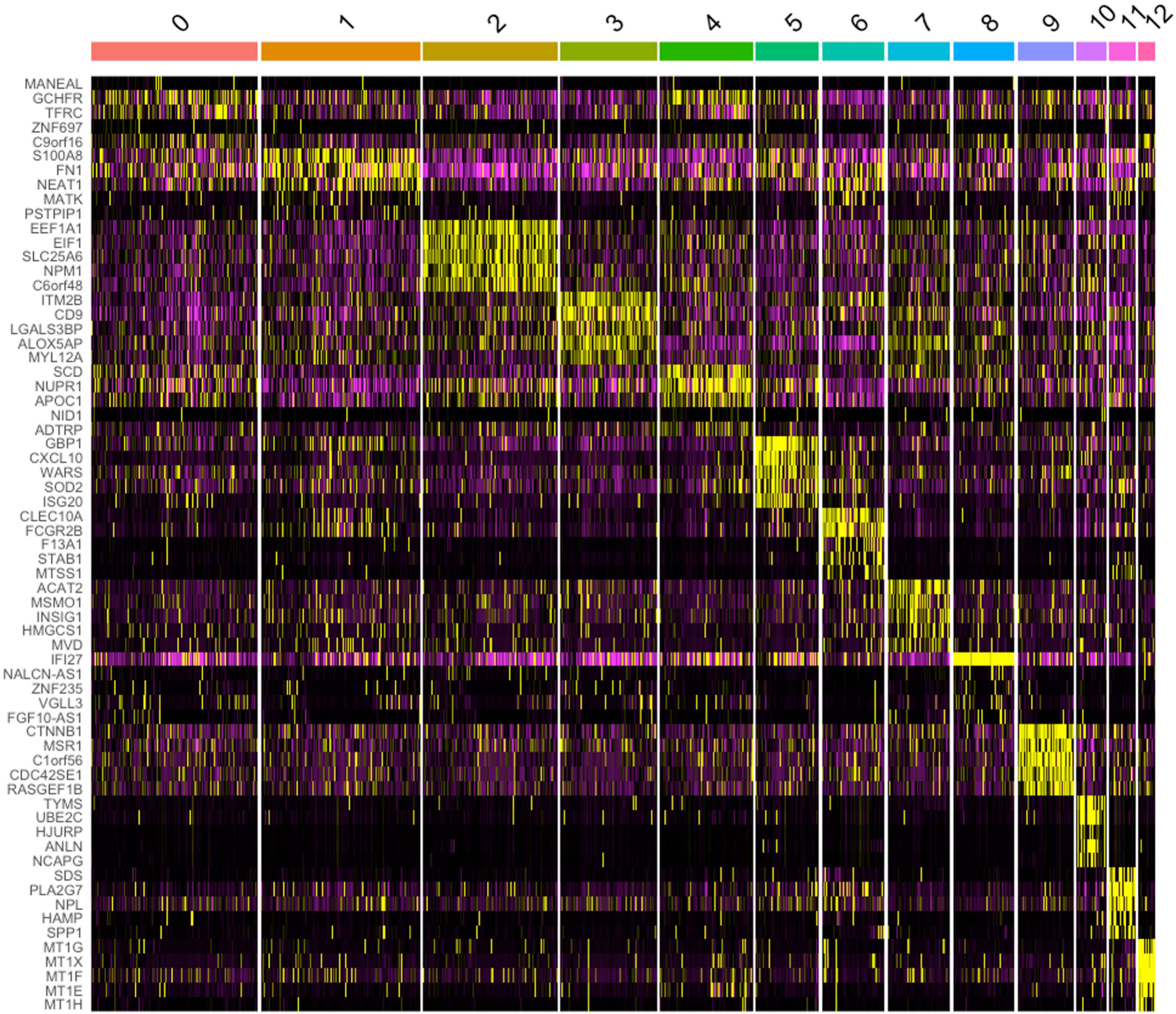

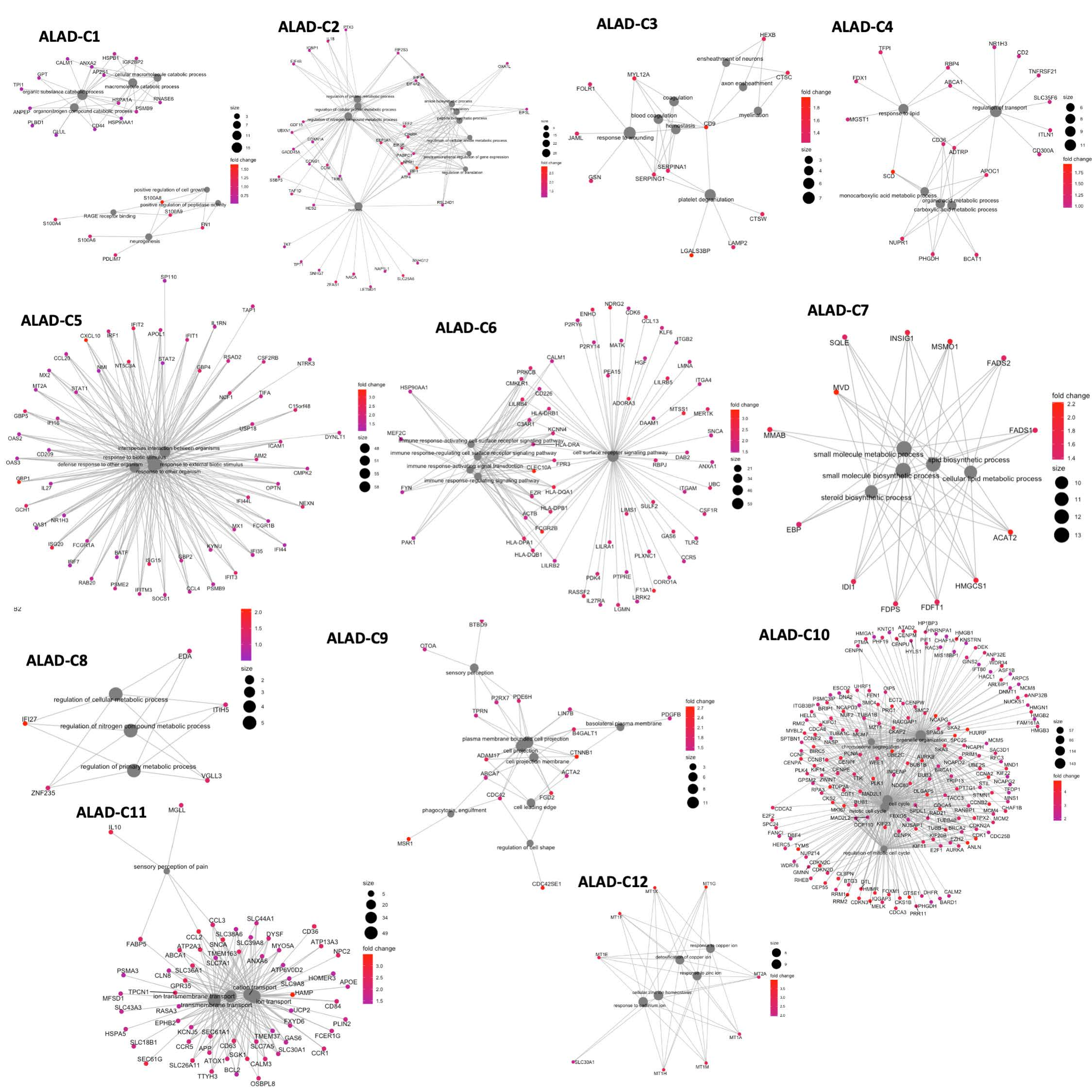

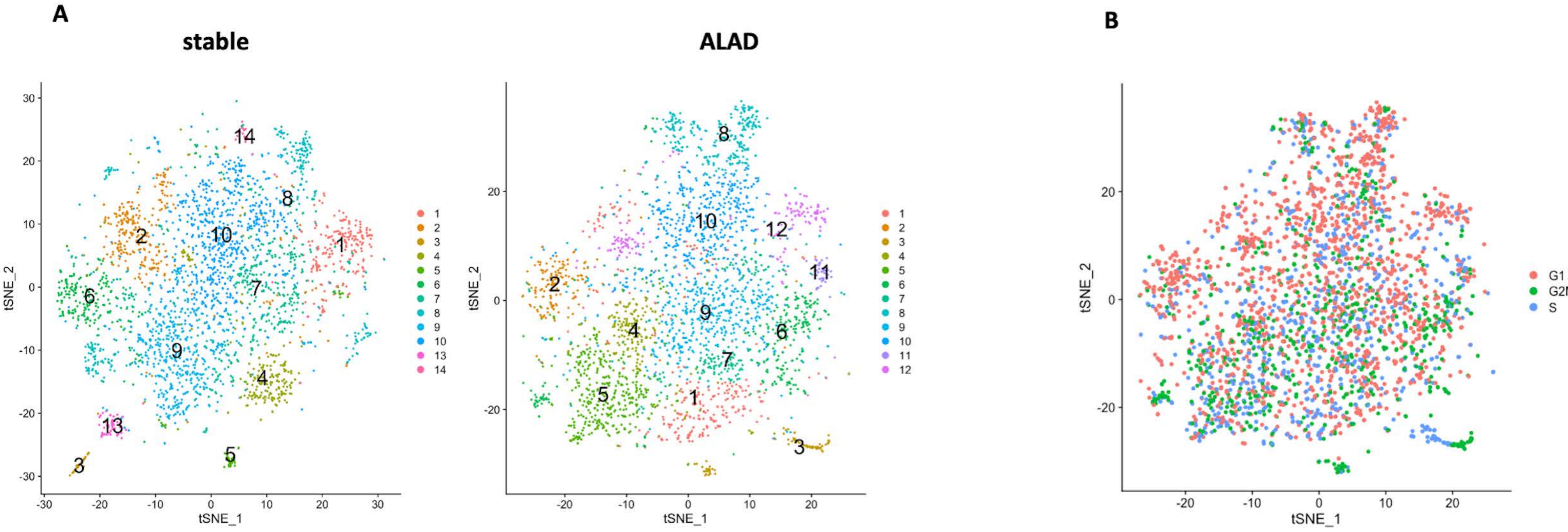

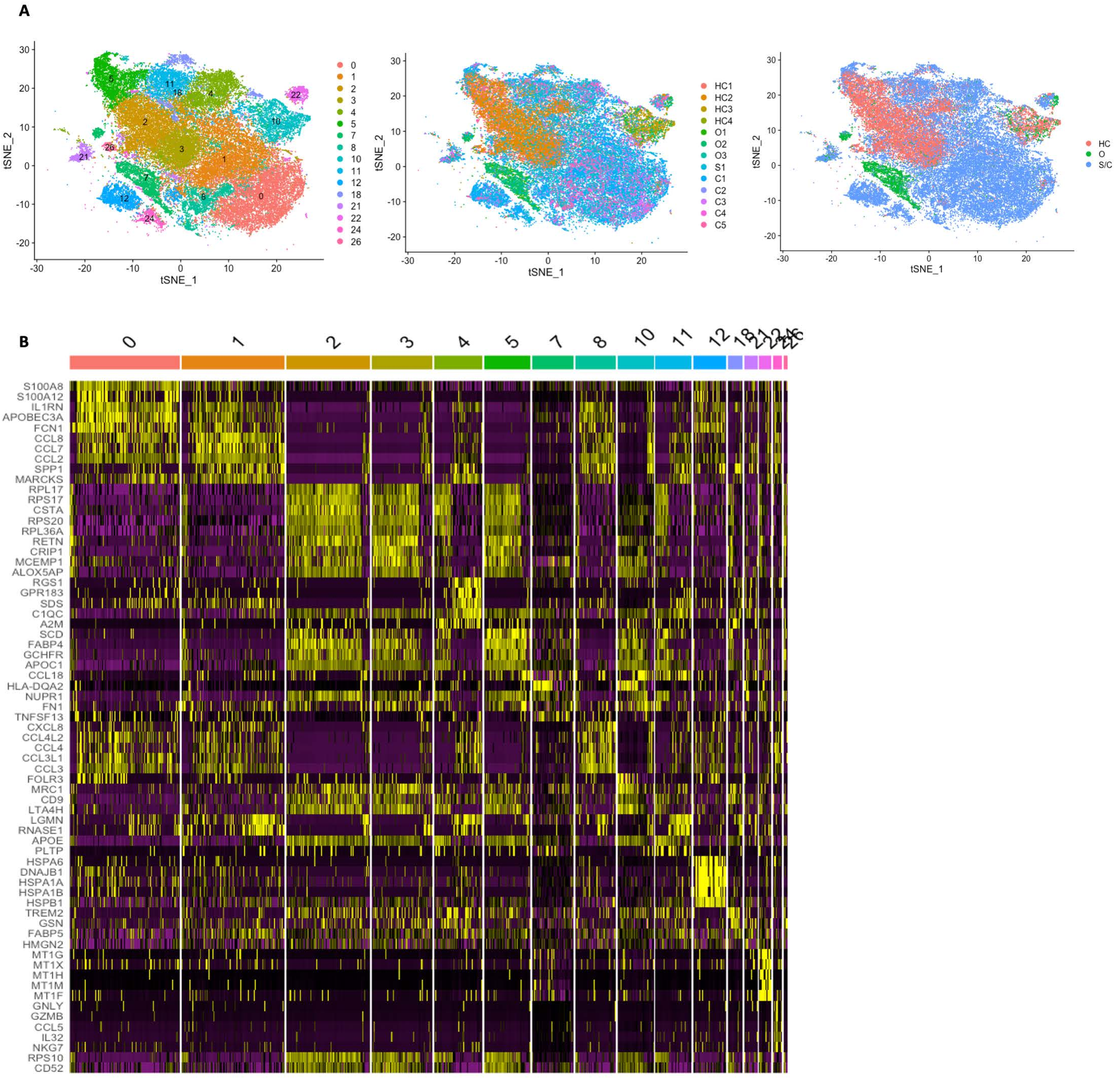

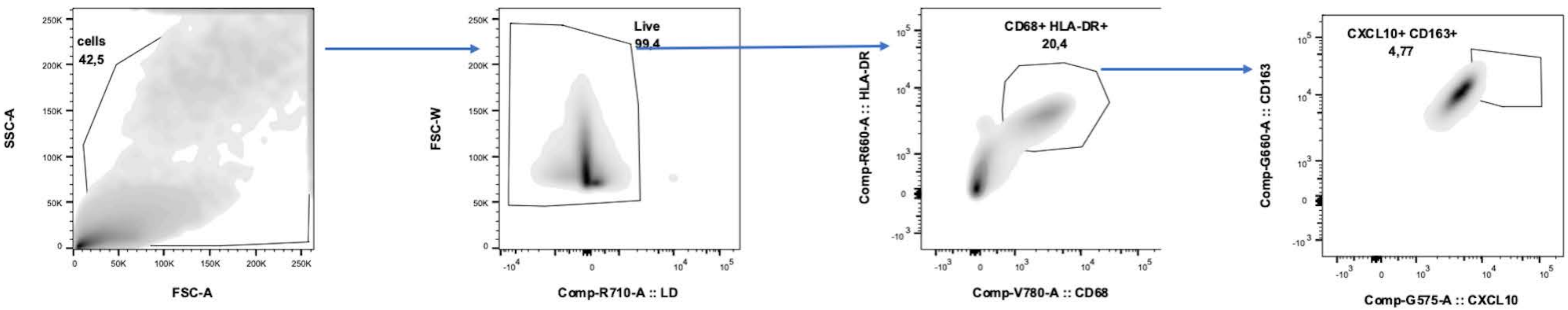
