## Supplemental Figure legend for "Pro-inflammatory alveolar macrophages associated with allograft dysfunction after lung transplantation"

**Figure S1. Visualization of quality control metrics for individual samples. A.** Violin plot demonstrating quality control of gene number (nFeature_RNA) between 200 and 5000, UMI count (nCount_RNA) between 1000 and 30,000 and mitochondrial gene percentage (percent.mt) below 10% of three stable and three ALAD BAL samples. **B.** Scatter plot showing linear relationship between gene number (Y axis) and UMI count (X axis) for each individual BAL sample.

**Figure S2. Integration cells from 3 stable and 3 ALAD BAL samples using SCTransform**. **A.** tSNE plot of integrated 3 stable BAL samples. **B.** tSNE plot of integrated 3 ALAD BAL samples.

**Figure S3. Expression of known macrophage transcripts across AM clusters.** Expression of CD68, CD163, MARCO, MRC1, CD11b, CD16, CD14, CD169, TGM2, CD86, TLR2, FTL, and a different HLA class II in AMs is shown (darker colour indicates higher relative expression).

**Figure S4. Signaling pathways that AM clusters communicate to each other.** Significant signaling pathways that are highlighted with red and larger circles have been used to identify potential functionality of AM clusters.

**Figure S5. Heatmap showing expression of top differentially-expressed genes in 13 AM clusters identified in AALAD AM samples.** Yellow indicates high expression.

**Figure S6. Pathway analysis on ALAD samples using GO and GSEA revealed distinct gene programs in the 13 clusters.**

**Figure S7. A.** tSNE plots showing stable (left) and ALAD (right) samples as outcome of Clustermap algorithm. In this tSNE format the colour and number represent the clusters with similar transcript signature. **B.** tSNE plot of AMs from ALAD with cell cycle stage indicated.

**Figure S8. A.** tSNE map pf AM clusters from healthy controls, mild and severe COVID-19 patients (data from Liao et al., 2020), showing AM cluster (left), origin of samples (middle), and COVID-19 status (right). **B.** Heatmap analysis of the top differentially-expressed genes in AMs identified by Liao et al. (2020). Cluster 0 shows similarity to the ISG cluster from ALAD patients, while cluster 22 shows similarity to the MT cluster in ALAD patients.

**Figure S9. Gating of AMs using flow cytometry.** Live, CD68+HLA-DR+ cells were identified as AMs. In this gate, CD163+CXCL10+ AMs could be identified.
